# Kinetic Analysis of Lipid Metabolism in Live Breast Cancer Cells via Nonlinear Optical Microscopy

**DOI:** 10.1101/827931

**Authors:** J. Hou, N. E. Reid, B. J. Tromberg, E. O. Potma

## Abstract

Investigating the behavior of breast cancer cells via reaction kinetics may help unravel the mechanisms that underlie metabolic changes in tumors. However, obtaining human *in vivo* kinetic data is challenging due to difficulties associated with measuring these parameters. Non-destructive methods of measuring lipid content in live cells, provide a novel approach to quantitatively model lipid synthesis and consumption. In this study, two-photon excited fluorescence (TPEF) was used to determine metabolic rates via the cell’s optical redox ratio (ORR) as reported by fluorescence intensity ratios of metabolic coenzymes, nicotinamide adenine dinucleotide (NADH) and flavin adenine dinucleotide (FAD+). Concurrently, coherent Raman scattering (CRS) microscopy was used to probe *de novo* intracellular lipid content. Combining non-linear optical microscopy and Michaelis-Menten-kinetics based simulations, we isolated fatty acid synthesis/consumption rates and elucidated effects of altered lipid metabolism in T47D breast cancer cells. When treated with 17β-Estradiol (E2), cancer cells showed a 3-fold increase in beta-oxidation rate as well as a 50% increase in cell proliferation rate. Similarly, the rate of *de novo* lipid synthesis in cancer cells treated with E2 was increased by 60%. Furthermore, we treated T47D cells with etomoxir (ETO) and observed that cancer cells treated with ETO exhibited a ∼70% reduction in β-oxidation. These results show the ability to probe lipid alterations in live cells with minimum interruption, to characterize both glucose and lipid metabolism in breast cancer cells via quantitative kinetic models and parameters.

**Statement of Significance:** Combining non-linear optical microscopy (NLOM) and deuterium labeling provides insight into lipid metabolism in live cancer cells during cancer development and progression. The dynamic metabolic data is modelled with Michaelis-Menten-kinetics to independently quantify the lipid synthesis and utilization in cancer cells. Changes in lipid levels are found to originate from *de novo* lipid synthesis using glucose as a source, lipid consumption from β-oxidation and lipid consumption from cell proliferation, processes that can separately analyzed with the Michaelis-Menten model. In this work, we isolate fatty acid synthesis/consumption rates and elucidated effects of altered lipid metabolism in T47D breast cancer cells in response to estradiol stimulation and etomoxir treatment, dynamic processes that cannot be easily observed without the application of appropriate models.

## 1. Introduction

The abnormal production of metabolites for the synthesis of cellular building blocks and signaling molecules is an emerging hallmark of cancer [1]. Cancer cells utilize the intermediate products of glycolysis to fuel the biosynthesis of amino acids, nucleotides and fatty acids to support fast cell proliferation. Different cancer cell lines exhibit varied glucose and lipid metabolic signatures correlated with their metastatic potential and cell proliferation rate both *in vitro* and *in vivo* [2-4]. Moreover, tumor metabolism has been shown to predict cancer cell response to chemotherapy in early stages [5]. Thus, quantitative analysis of cancer cell metabolic kinetics is of great importance in characterizing cancer cell behavior and unraveling the role of cell metabolism in cancer progression and transformation.

Biochemically, cell metabolism is achieved through multiple series of reactions of varying complexities, catalyzed by proteins or catalytic RNAs. The Michaelis-Menten equation is often used to quantify enzyme related metabolic behavior in normal and mutated cells (equation 1).

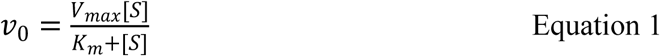

where, V_max_ is the maximum rate achieved by the reaction, [S] is the substrate concentration, and K_m_ is the Michaelis constant or substrate concentration at which the reaction rate is half of V_max_. By combining fluorescence, isotope and radio labeling with Michaelis-Menten kinetics, cell metabolic rates can be investigated quantitatively. In a clinical study, the production of tritiated water was assessed in human patient cells treated with [1β-3H(N)]-androstenedione [6]. The cells that expressed either wild-type CYP191A genes or its corresponding mutations (Y81CL451P) were observed to have different tritiated water production rate. By fitting with Michaelis-Menten kinetics, V_max_, K_m_ and V_max_ / K_m_ (catalytic efficiency) were calculated for each group and were observed to be affected by aromatase deficiency. Similarly, Michalis-Menten parameters were employed to study intracellular hydrolysis for the detection of breast cancer by measuring intracellular fluorescein intensity [7]. However, most of these traditional labelling methods relied on exogenous chemicals, which may compromise the living biological system in unknown ways. Moreover, the requirement for complicated sample preparation and time-consuming data acquisition impede the possibility of dynamic studies with high spatial and temporal resolution.

Multi-modal, non-linear optical microscopy (NLOM) enables minimally interrupted evaluation of cell metabolism, with high spatial and temporal resolution. Previously, two-photon excited fluorescent (TPEF) microscopy was used to monitor cell glucose metabolism over a range of oxygen consumption conditions relevant for cancer imaging [8]. By studying the intra-cellular lipid droplets with both coherent Raman scattering (CRS) microscopy and spontaneous Raman spectroscopy [9], Shuhua et al. reported increased cholesterol accumulation in prostate cancer cells that correlated with cancer aggressiveness. Recently, CRS was utilized with deuterium labelling to quantify lipid metabolism. Zhang *et al.* studied cancer anabolism after epithelial-mesenchymal transition by tracking the deuterium and alkyne vibrational signal with CRS [10]. These research findings underline the potential of utilizing NLOM to quantify live cell metabolism with high spatial and temporal resolution.

In this study, quantitative models were derived and used to describe alterations of the glucose metabolic pathways in the context of lipid production and consumption in cancer cells. We use TPEF and CRS microscopy to determine glucose metabolic rates and lipid content, respectively. In addition, deuterium labeling was utilized to investigate *de novo* lipid synthesis in cancer cells. We monitored fatty acid synthesis from deuterated glucose and subsequent consumption by tracking the carbon-deuterium signal in a pulse chase experiment. Recently, Luyuan *et al.* observed changes of glucose-supplied lipid renewal in sebaceous gland over qualitatively by combining the CRS with deuterium labelling [11]. However, it was still based on observations of total lipid content changes and cannot quantify lipid synthesis and consumption independently. Here, we combined CRS, deuterium labeling and Michaelis-Menten-kinetics modeling and isolated fatty acid synthesis and consumption rates. We revealed the effects of altered lipid metabolism in the estrogen receptor positive (ER+) T47D breast cancer cells when treated with 17ß-Estradiol (E2). We find that the T47D breast cancer cells exhibited elevated levels of cell proliferation in addition to increased lipid synthesis and consumption rates when activated by E2. The high lipid *de novo* synthesis rate corroborated the rise of the glycolytic rate measured by TPEF. We also treated T47D breasts cancer cells with etomoxir (ETO) in order to investigate the modeling efficiency in extreme beta-oxidation inhibition conditions.

## 2. Material and Methods

### 2.1. Model Development

Lipids, such as triglycerides and phospholids, are generated from glucose-derived glycerol and mitochondrial derived fatty acids. Fatty acid synthesis takes place in the cytosol of the cell, where mitochondrial citrate serves as a precursor to palmitate, which is converted into other fatty acids.

In order to develop a mathematical model to describe lipid metabolism, the rate-limiting step of the process must be assumed. In chemical kinetics, the overall rate of a reaction is often approximated by the slowest step, known as the rate-limiting or rate determining step (RDS). Fatty acid synthesis is regulated by acetyl-CoA carboxylase (ACC), which is activated by citrate and inhibited by fatty acid palmitate [12]. On the other hand, fatty acid oxidation is regulated by malonyl-CoA, which inhibits carnitatin palmitoyl transferase (CPT1) activity. CPT1 is overexposed in many cancer cells and is believed to be responsible for suppression and reduction of cell growth in vitro [13].

Assuming the rate of lipid synthesis and consumption is dependent on the rate-limiting enzymes described above, kinetic models can now be established. A similar approach was taken in the development of a complete pathway model for lipids in *Escherichia coli* biosynthesis [14]. Equations 2 and 3 show single substrate Michaelis-Menten kinetics with and without inhibition, respectively, where [P] is the product concentration, [S] is the substrate concentration, [E] is the enzyme concentration, k_cat_ is the rate of catalysis, K_m_ is the Michaelis-Menten constant and K_i_ is the inhibition rate constant [7].

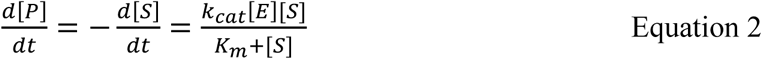

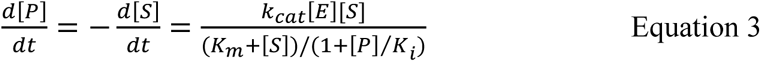

Experimental data in this study provided intracellular lipid content stemming from deuterated glucose. This implies that the maximum lipid content (L_max_) was less than the number of total initial glucose molecules (G_0_). Applying Equation 2 to describe the case of lipid consumption (beta-oxidation), we obtain:

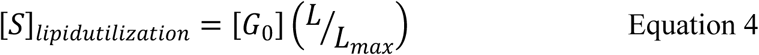

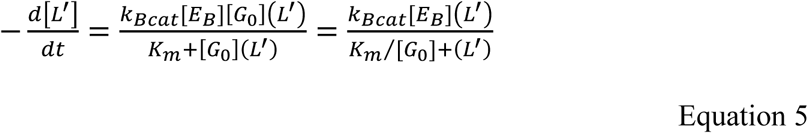

where k_BCat_ is the catalytic rate of lipid beta-oxidation (regulated by malonyl-CoA), [E_B_] is the concentration of malonyl-CoA, and L’ = L/L_max_ is the normalized lipid content. In similar metabolic studies, values of K_m_ range from 0.00019 – 0.82mmol/L [14]. In this study, [G_0_] = 17 mmol/L. Therefore, 1.12 × 10^−5 ≤^ K_m_/[G_0_] ≤ 0.048 mmol/L and the resulting denominator reduces to L’. We thus obtain:

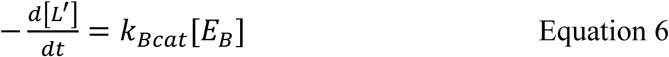

Lipid consumption is also affected by cell proliferation [15]. Thus, we applied a first order lipid consumption rate (K_p_) to equations 6 to develop a more inclusive rate model by taking the rate of lipid consumption by cell proliferation into account. This results in a sum of a zero order and a lipid dependent first order equation:

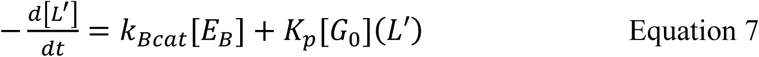

Similarly, equations 6 and 7 can be adjusted to describe the rate of *de novo* lipid synthesis with or without cell proliferation:

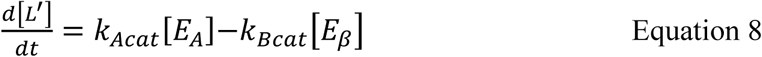

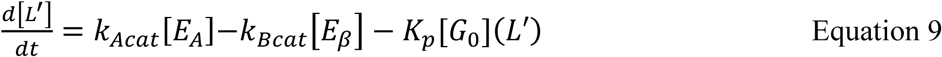

where k_ACat_ is the catalytic rate of lipid synthesis (regulated by acetyl-CoA carboxylase) and [E_A_] is the concentration of acetyl-CoA carboxylase. The models included in this study are summarized in table 1. They were fitted and tested against lipid metabolism data gathered using non-linear optical microscopy. Kinetic parameters were calculated from non-linear regression of the Michalis-Menten type equation by Matlab software. The goal is to quantitatively correlate kinetic parameters to internal and external conditions in T47D cancer cells.

**Table 1.**
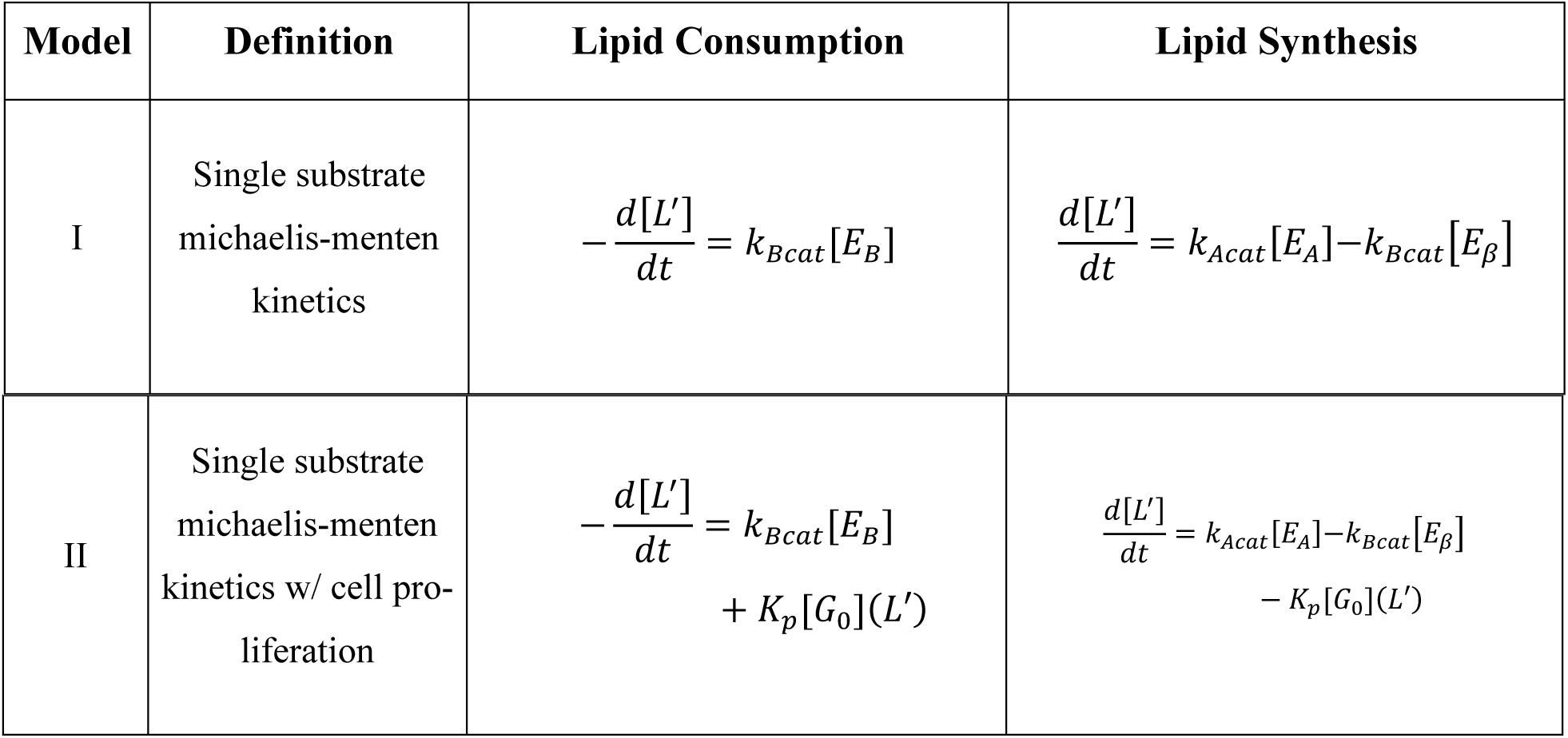
Models tested in our study.

### 2.2. Matlab simulation

Data from lipid synthesis and pulse chase experiments were normalized based on the maximum lipid content observed in each experiment (L’=L/L_max_). Then the data was fitted to four models using a Matlab minimization routine, ‘fminsearch’. First, using a numerical ordinary differential equation solver (ODE45), the code takes the measured deuterated lipid data and returns a predicted fitting. sSecondly, the code computes the sum of the square of differences between the predicted and experiment data. The iterative process continued until the algorithm has found the fitted parameters that minimize this difference. Accuracy of fitting was measured using the square of Pearson’s correlation coefficient (r^2^). The correlation coefficient indicates how well the calculated curve fits the original data, with a value ranging from zero to one. The closer to 1.0, the better the fit. Models were eliminated if r^2^ values were low (r_2_ < 0.90) or fitted parameters resulted in a negative value (not physically possible).

### 2.3. Nonlinear Optical Microscopy

A 76MHz Nd:Vanadate laser at 1064nm (picoTrain, High-Q) was used, producing a 7ps pulsed laser beam (Stokes beam). The second harmonic of the same laser (532nm) was used to pump an optical parametric oscillator (OPO; Levante, Emerald OPO, Applied Physics & Electronics Inc.) to provide the corresponding pump beam for imaging either the C-H lipid distribution (817nm for a Raman shift of 2845 cm^-1^) or C-D lipid signal (864nm for a Raman shift of 2175 cm^-1^). The Stokes beam was modulated with an acousto-optic modulator (12465, Crystal Technology, Inc.) at 10MHz. Both the Stokes beam and pump beam were spatially and temporally overlapped and delivered to a laser-scanning microscope (IX71, Olympus). The CRS signal was detected in the forward direction. For this purpose, the pump beam was focused onto a photodiode (FDS1010, Thorlabs, Inc.) with a high O.D. bandpass filter for blocking the Stokes beam. The output of the photodiode was filtered with an electronic bandpass filter (BBP-10.7+, MiniCircuits, Inc.) and demodulated by a homemade lock-in amplifier to detect the Raman loss signal. The cells were imaged by a 60X water objective (1.2NA, UPlanSApo, Olympus) and the CRS signal was collected in the forward direction by a condenser (NA = 1.05). An olive oil sample was measured before each experiment for calibration purposes.

The optical redox ratio was measured on a Zeiss laser-scanning microscope (LSM510) equipped with a femtosecond laser source (Chameleon, Coherent Inc.). The two-photon excited fluorescence signal of NADH (740nm excitation and 480nm±50nm emission filter) and FAD+ (900nm excitation and 530nm±50nm emission filter) was generated sequentially. The optical redox ratio was then calculated as

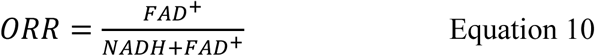

The cells were imaged through a 40X water objective (0.8NA, LUMPlanFl, Olympus) and the fluorescence signals were collected in the epi-direction. A freshly-prepared fluorescein solution (0.02 μM at pH=7) was imaged before each measurement session as a reference.

### 2.4. Cell Culture

The T47D breast cancer cell line was acquired from ATCC. The normal cell culture medium is made from Advance DMEM/F-12 culture medium (A2494301, ThermoFisher scientific) supplemented with 5% fetal bovine serum (12676011, Corning), 1% glutaMAX (35050-061, Life Technologies), 147.5 mg/L l-arginine (A5006, Sigma-Aldrich), 91.25 mg/L l-lysine (L5501, Sigma-Aldrich) and 3151 mg/L normal glucose (G7021, Sigma-Aldrich). The deuterated medium was prepared by replacing the normal glucose with deuterated glucose (552003, Sigma-Aldrich) of the same concentration. The cells were cultured at 37 °C and 5% CO_2_. The cells were monitored daily and the culture medium was changed every two to three days. For all the experiments, the cells were detached and collected at around 60% confluence with TrypLE (12604013, Life Technologies).

For lipid synthesis experiments, the cells were harvested and plated onto coverslips at 50,000 cells/well in 24 well plates. The cells were incubated in normal culture medium for 24 hours to allow the cells to recover. The cells were then switched to serum free culture medium overnight for cell cycle synchronization. Subsequently, the cells were exposed to deuterated culture medium (considered T=0). One plate of the cells was treated with 10^−8^ M 17β-estradiol (E8875, Sigma-Aldrich) and the other plate of the cells was treated with DMSO at the same concentration. We fixed two wells (two replicates) for each plate at T = 0h, 2h, 3h, 4h, 5h, 6h, 9h, 12h in 4% formaldehyde.

For pulse chase experiment, the cells were harvested and plated onto coverslips at 50,000 cells/well in 24 well plates. The cells were incubated in normal culture medium for 24 hours to allow the cells to recover and switch to deuterated glucose medium for 24 hours. To test the effect of 17β-estradiol (E2), one plate of the cells was treated with 10^−8^ M 17β-estradiol and the other plate of cells was treated with 10^−8^ M DMSO. To test the effect of etomoxir, one plate of the cells was treated with 4*10^−5^ M etomoxir and the other plate of the cells was treated with 4*10^−5^ M vehicle control. The cells were then switched to normal glucose medium (considered T=0) and we fixed two wells (replicates) of the control group and experimental group at times T=0h, 2h, 4h, 6h, 9h, 12h.

### 2.5. Lipid Quantification

The CRS images were imported into imageJ. The image intensities were first calibrated by the olive oil reference sample as,

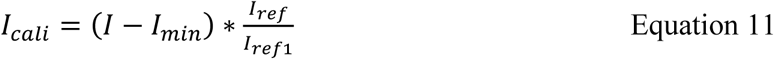

where *I*_*call*_ is the calibrated signal, *I* is the direct measured signal intensity, *I*_*min*_ is the minimum intensity in the image,*I*_*ref*_ is the mean intensity of the olive oil sample measured before the experiment and *I*_*ref*1_ is the mean intensity of the reference sample for the first experiment.

In each field of view, both the normal lipid distribution and deuterated lipid distribution were taken sequentially by tuning the wavelength of pump beam. All the images were first imported into Matlab to calibrate their intensities and remove the background by Otsu thresholding. Then, the processed images were analyzed in imageJ where 25 lipid droplets were randomly selected and manually outlined. The average normal signal and deuterated signal intensities within each lipid droplet was measured. The lipid content was defined as the intensity ratios of the deuterated lipid and normal lipid of the same droplets.

## 3. Results and Discussion

### 3.1 Evaluating the efficiency of using different Michaelis-Menten model to simulate cancer metabolism

To validate the modeling approach, we measured the changes of deuterated lipid signal in T47D breast cancer cells treated with either E2 or DMSO. T47D is an estrogen receptor positive breast cancer cell line. Based on previous publications, ER+ breast cancer cells were sensitive to E2 treatment and responded with altered cell metabolism and proliferation. We used this model to test the sensitivity and accuracy of our simulation to clinically related cell metabolic changes.

We first cultured the cells in deuterated glucose medium for 24 hours to label all intracellular lipid content with deuterium. Then, we switched all the cells to normal glucose culture medium and started to monitor the intensity changes of the coherent Raman signal that originated from the carbon-deuterium vibrational mode. The pulse chase experiment lasted for 24 hours and the decay of deuterated lipid signal in T47D cells treated with either DMSO or E2 were recorded and fitted with single substrate Michaelis-Menten kinetics (equation 6), and single substrate Michaelis-Menten kinetics with cell proliferation (equation 7).

The simulation results are summarized in Table 2 and the goodness of fit for each model was evaluated with an r^2^ value. From the table, reduced single substrate Michaelis-Menten kinetics with cell proliferation (Model II) provided the best fit. Thus, we utilized model II for all of the following experiments. Note that similar Michaelis-Menten parameter values were achieved for metabolite formation in other breast cancer studies [16].

**Table 2.**
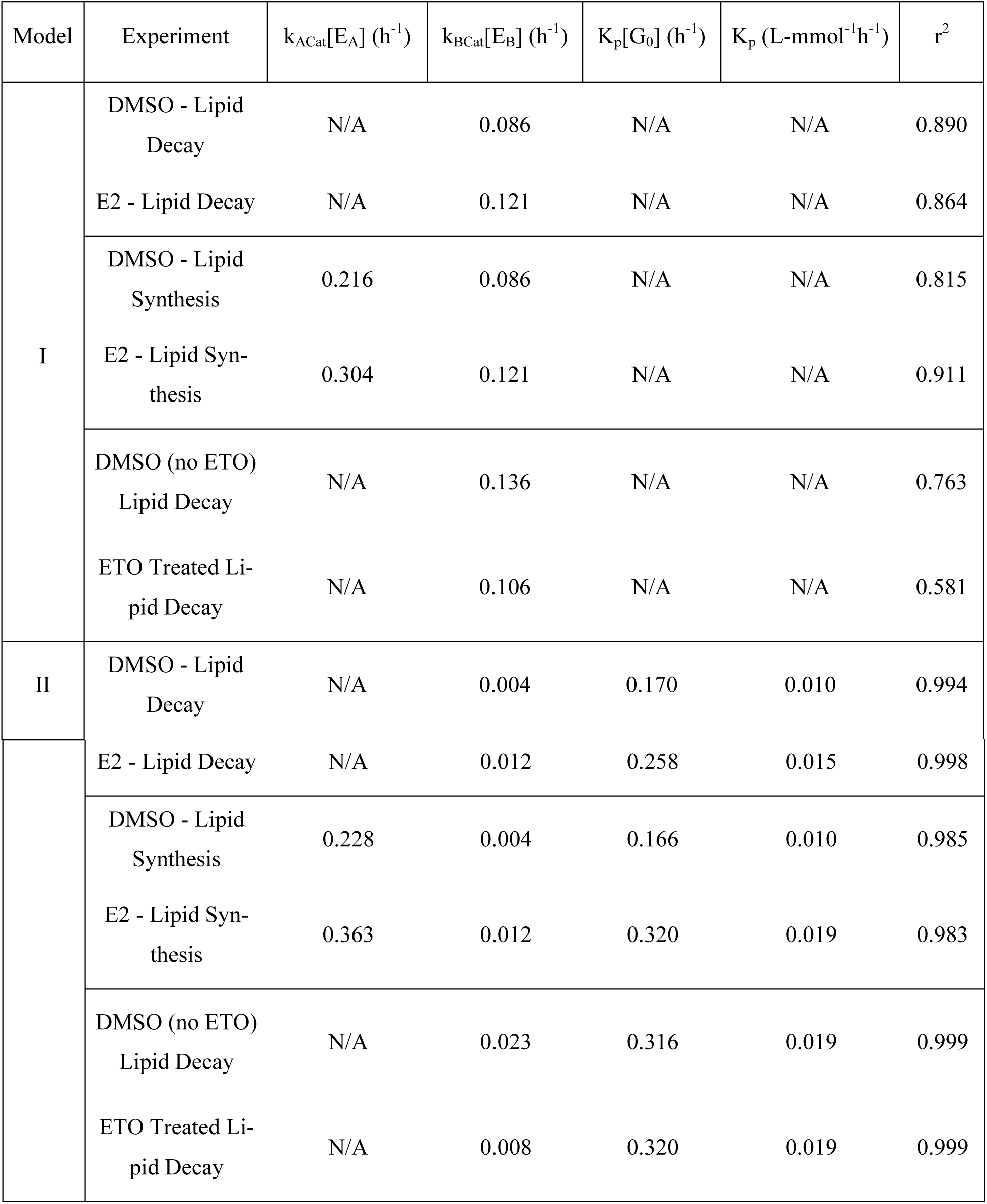
Simulation results of T47D cancer cells treated with DMSO or E2.

| Model | Experiment | $k_{ACat}[E_A] \text{ (h}^{-1}\text{)}$ | $k_{BCat}[E_B] \text{ (h}^{-1}\text{)}$ | $K_p[G_0] \text{ (h}^{-1}\text{)}$ | $K_p \text{ (L-mmol}^{-1}\text{h}^{-1}\text{)}$ | $r^2$ |
| --- | --- | --- | --- | --- | --- | --- |
| I | DMSO - Lipid Decay | N/A | 0.086 | N/A | N/A | 0.890 |
|  | E2 - Lipid Decay | N/A | 0.121 | N/A | N/A | 0.864 |
|  | DMSO - Lipid Synthesis | 0.216 | 0.086 | N/A | N/A | 0.815 |
|  | E2 - Lipid Synthesis | 0.304 | 0.121 | N/A | N/A | 0.911 |
|  | DMSO (no ETO) Lipid Decay | N/A | 0.136 | N/A | N/A | 0.763 |
|  | ETO Treated Lipid Decay | N/A | 0.106 | N/A | N/A | 0.581 |
| II | DMSO - Lipid Decay | N/A | 0.004 | 0.170 | 0.010 | 0.994 |
|  | E2 - Lipid Decay | N/A | 0.012 | 0.258 | 0.015 | 0.998 |
|  | DMSO - Lipid Synthesis | 0.228 | 0.004 | 0.166 | 0.010 | 0.985 |
|  | E2 - Lipid Synthesis | 0.363 | 0.012 | 0.320 | 0.019 | 0.983 |
|  | DMSO (no ETO) Lipid Decay | N/A | 0.023 | 0.316 | 0.019 | 0.999 |
|  | ETO Treated Lipid Decay | N/A | 0.008 | 0.320 | 0.019 | 0.999 |

### 3.2 Kinetic model reveals the effect of 17β-estradiol on an ER positive breast cancer cell line

Previously, increased cell proliferation rate and cancer invasiveness was observed in T47D cancer cells upon treatment with E2 [17]. The change of cancer cell phenotype was known to alter cancer cell lipid consumption and lipid synthesis. To uncouple lipid synthesis from consumption, both the deuterated lipid decay and deuterated lipid accumulation needed to be monitored. Thus, we studied 1) the increase of deuterated signal in normal T47D cells cultured in deuterated glucose medium and 2) the decrease of deuterated signal in fully deuterated T47D cells cultured in normal glucose medium. The dynamic changes of T47D cells treated with either E2 or DMSO were monitored for 24 hours. The deuterium signal decay was first fitted with equation 7 to get the lipid oxidation rate (k_BCat_[E_B_]) and cell proliferation (K_p_[G_0_]). Then the lipid accumulation data were fitted with equation 9 to find the lipid synthesize rate, k_ACat_[E_A_] where k_BCat_[E_B_] and K_p_[G_0_] were directly derived from the corresponding lipid consumption pulse chase experiment.

The CRS images taken at 2175 cm^-1^ showed significant deuterated lipid deposition in cancer cells cultured with deuterated glucose after 24 hours of culturing in deuterated medium (fig 1a and b). After switching to the normal glucose culture medium, the deuterated lipid droplets signal intensity dropped to background levels within 12 hours in cancer cells treated with E2 (fig. 1c). At the same time point, the signal intensity was 110 times higher in the cancer cells treated with DMSO. In the lipid synthesis study, the deuterated lipids became visible as early as 2 hours in both conditions. Both cancer cells in the DMSO and E2 treated groups demonstrated a rapid increase in deuterated signal from 2 hours to 6 hours (fig. 1d). After 9 hours of culturing, the deuterated signal reached a plateau in both conditions. However, the cells treated with E2 demonstrated 1.6 times higher signal intensity over the control group after reaching the equilibrium state.

**Figure 1.**
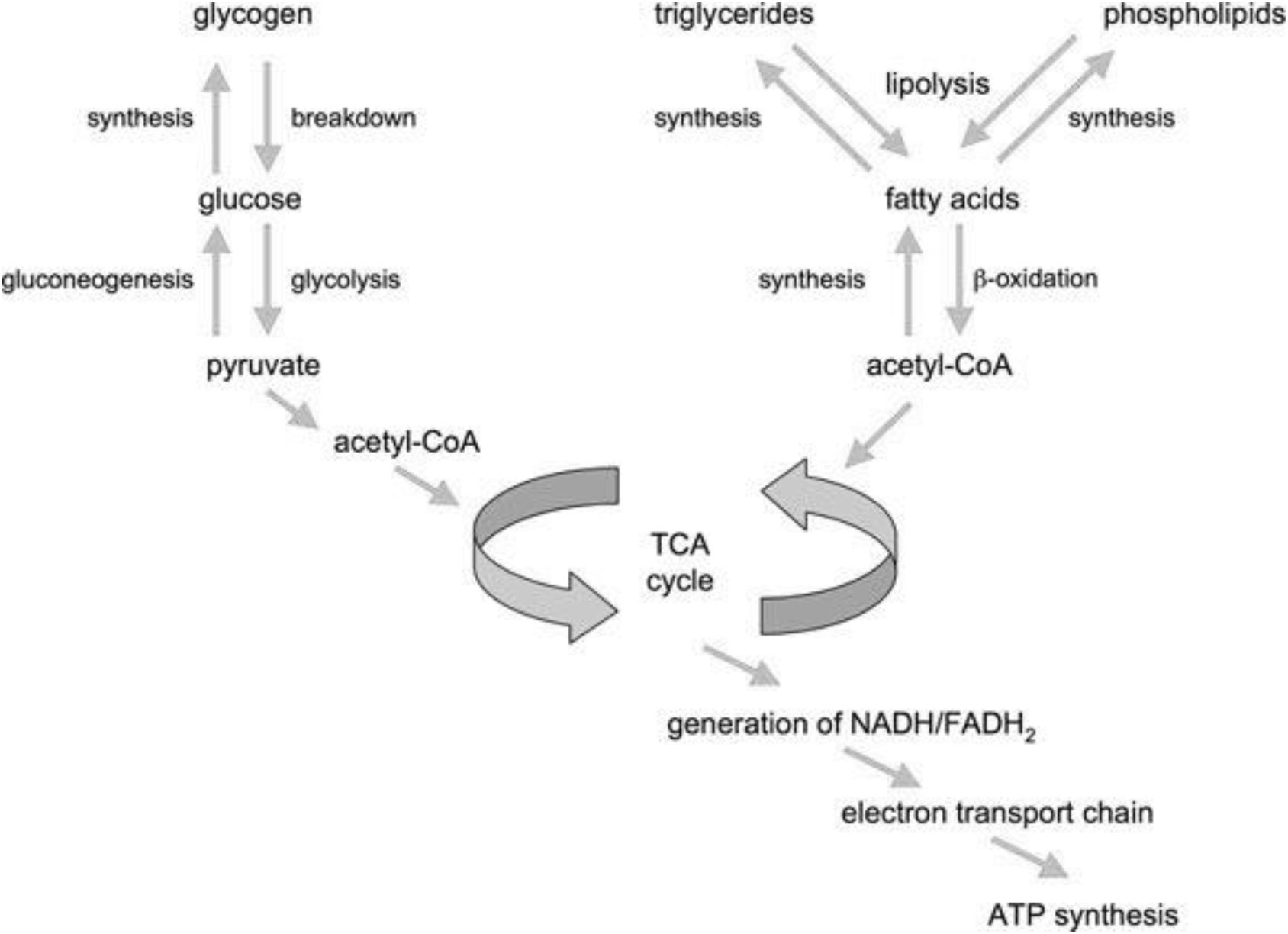
Overview of fat and sugar synthesis and breakdown pathways.

**Figure 2.**
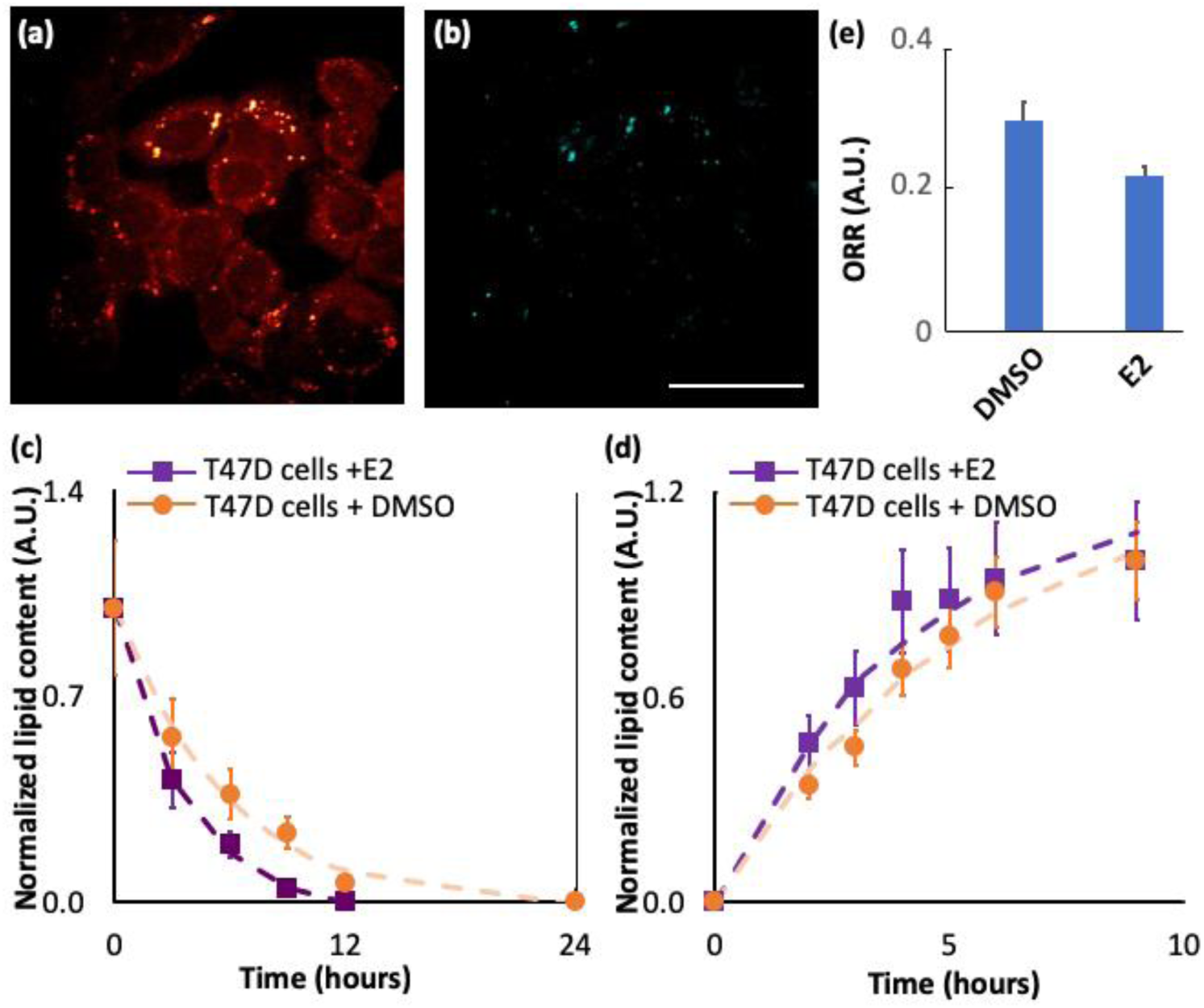
Deuterated lipid pulse chase experiments of lipid synthesis and lipid consumption. Representative (a) normal lipid distribution (817nm pump laser) and (b) corresponding deuterated lipid distribution (869nm pump laser) in T47D cancer cells at time = 0 hour. (c) Measured decay of normalized lipid content in T47D cells (control) and17β-Estradiol (E2) treated T47D cells, fitted to a modified single substrate lipid consumption with cell proliferation kinetic model. (d) Measured transient of lipid synthesis in T47D cells (control) and17 E2 treated T47D cells, fitted to a modified single substrate lipid synthesis with lipid consumption and cell proliferation kinetic model. (e) The optical redox ratio (ORR) for T47D cells treated with either DMSO or E2. Error bars represent the standard deviation from 25 randomly picked lipid droplets. Scale bar 20um.

From the NLOM images, the signal intensities in cancer cells treated with E2 demonstrated higher decay rate indicating increased lipid consumption. By fitting the data to our model, we found that the increased lipid consumption was due to enhanced β-oxidation rate and cell proliferation. On average, we found about 200% increase of the k_BCat_[E_B_] value (β-oxidation rate) and 50% increase of the K_p_[G_0_] value (cell proliferation) (table 3). Our quantitative data retrieved from NLOM imaging and mathematical modeling matches with previous publications. Using colorimetric assay, Lai and *et al.* reported that ER+ cancer cells treated with 10^−8^ M E2 grew 43% faster [18]. In several independent studies, the activation of ER by estradiol has also been reported to upregulate the expression of trifunctional protein β-subunit (HADHB) and fatty acid elongation of very long-chain fatty acids 2 (ELOVL2) to increase β-oxidation [19, 20].

**Table 3.**
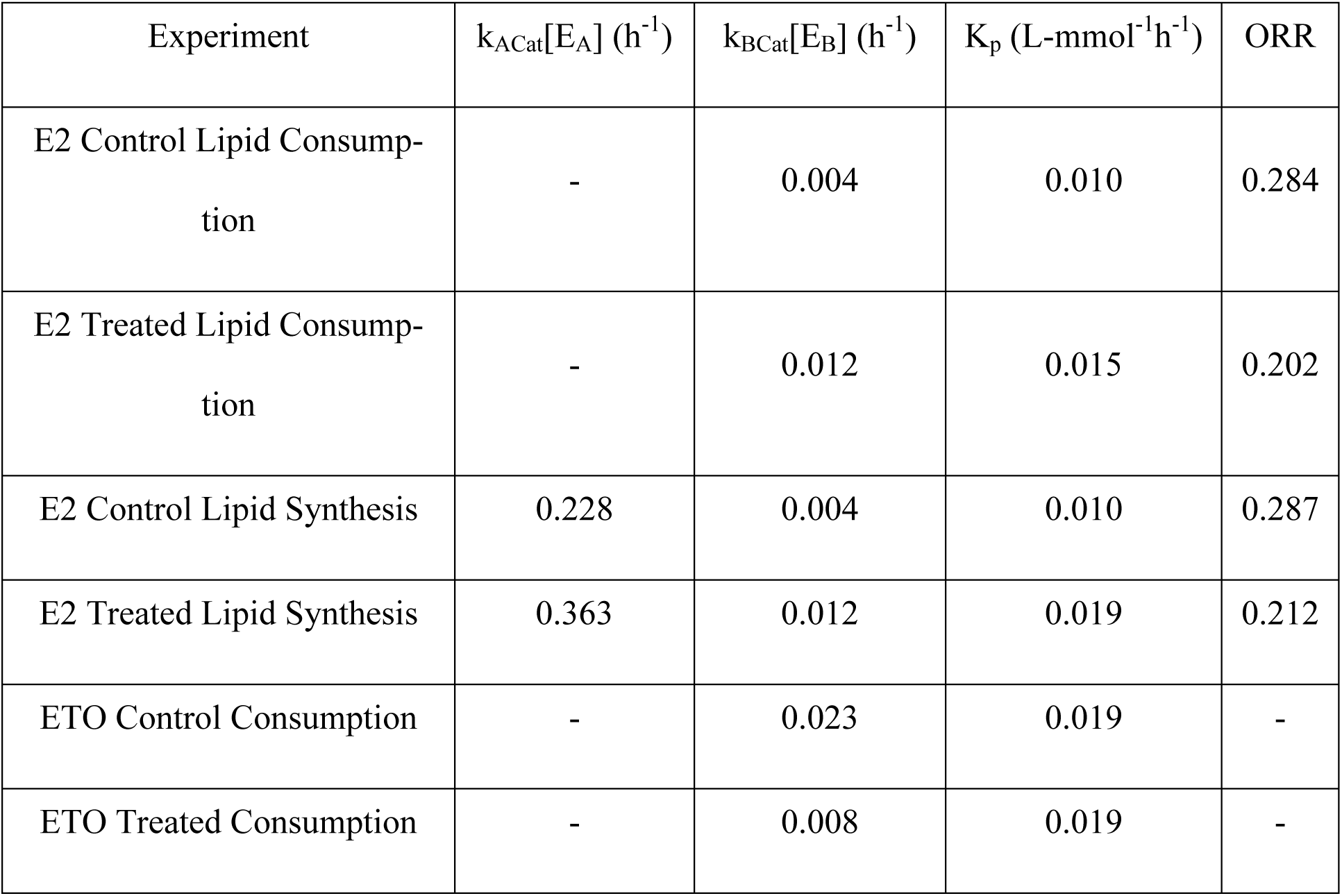
Summary of kinetic parameters based on a Michaelis-Menten type model describing lipid synthesis and consumption via beta-oxidation and cell proliferation.

Interestingly, our lipid synthesis study demonstrated that the cancer cell treated with E2 display increased *de novo* lipid synthesis along with enhanced lipid consumption. The E2 treated cancer cells had 40% higher k_ACat_[E_A_] values compared to the cells treated with DMSO. To further validate our observations, we measured the ORR of T47D cells under both conditions. Previously, we reported that a decrease of ORR is associated with a higher glycolytic rate, which can promote biosynthesis by providing precursors through aerobic glycolysis [8]. Here, T47D cells exhibited a 30% decrease in ORR when treated with E2 suggesting an increase of lipid synthesis as predicted by our model.

Biologically, the treatment of estradiol will stimulate estrogen receptor positive breast cancer cells’ proliferation and motility. The altered cancer phenotype imposed an increased demand on ATPs and biomass, which is supported by an increase in lipid β-oxidation and *de novo* lipid synthesis. Our NLOM imaging method and mathematical modeling can quantitatively monitor the change of the metabolic rate, thus providing a powerful tool for cell metabolism studies.

### 3.3 Sensitivity of the model to changes of lipid metabolism

To further test the utility of the kinetic model in extreme conditions, breast cancer cells were treated with etomoxir (ETO). ETO constrains cell β-oxidation by inhibiting carnitine palmitoylytransferase-1 (CPT-1) on the mitochondria membrane. Similar to above experiments, T47D breast cancer cells were first cultured in deuterated glucose for 24 hours before switching to normal culture medium. The deuterated signal decay from the lipid droplets was measured over 12 hours.

At time T=0, the signal intensity was similar in both experimental groups. In the control group, the deuterium signal dropped to background levels after 9 hours of culturing (fig. 3a and b). However, the deuterated lipid droplets were still visible in the T47D cells treated with ETO after 12 hours and remained at a constant level (fig. 3 c-e). This remaining lipid content mostly originated from inactive cells. It is known that ETO treatment adversely affects normal metabolic pathways and cell viability [21]. Thus, the lipid metabolism was no longer mediated by enzymatic reactions and pure Michaelis-Menten analysis was inadequate under such conditions. To account for this limitation, we subtracted all the measured data points by the lipid content which remained constant after 9 hours to effectively eliminate the lipid content from inactive cells. With the adjusted model, the ETO treated cells demonstrated a 65% reduction in β-oxidation as the coefficient k_BCat_[E_B_] dropped from 0.023 mmol/(L•h) to 0.0008 mmol/(L•h). Moreover, the cell proliferation rate remained unchanged between the two groups. It can be expected that the ETO treatment greatly reduces the cell β-oxidation rate but has little effect on cell proliferation in the cancer cells that can tolerate ETO treatment. This experiment demonstrated the limitation of Michaelis-Menten modelling in that that it can only be used for describing enzymatic processes. When analyzing such extreme conditions, the inactive cells must be excluded to ensure the accuracy of the modelling.

**Figure 3.**
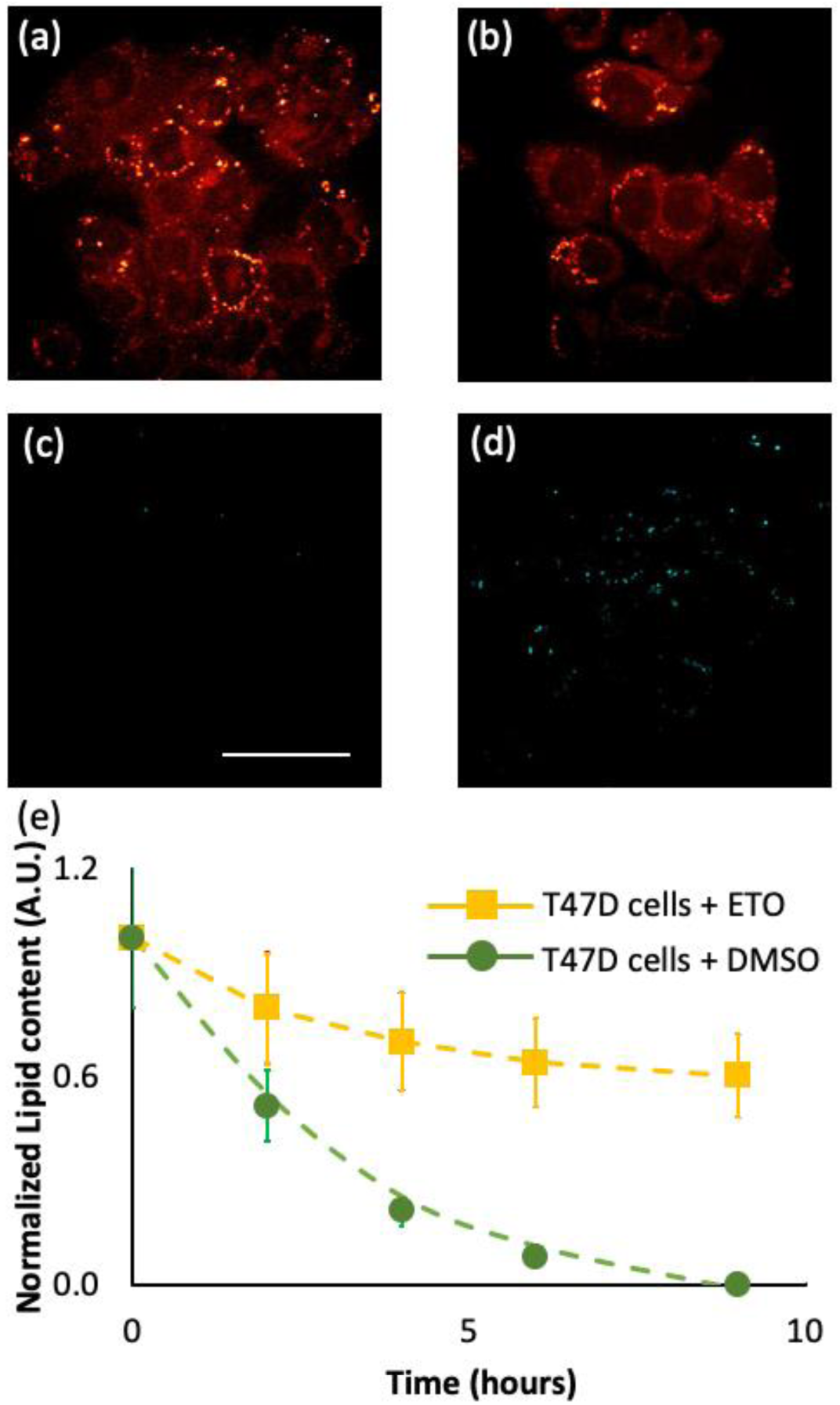
Pulse chase experiments of ETO treated T47D cancer cells. CRS images of representative normal lipid distribution at T=12 hours for (a) ETO treated T47D cells and (b) DMSO treated T47D cells. CRS images of representative deuterated lipid distribution at T=12 hours for (c) ETO treated T47D cells and (d) DMSO treated T47D cells. (e) Measured decay of normalized lipid content in T47D cells treated with either DMSO or ETO, fitted to a modified single substrate lipid consumption with cell proliferation kinetic model. The error bars represent the standard deviation from 25 randomly picked lipid droplets.

## 4. Conclusion

We have combined CRS imaging with quantitative kinetic modeling to reveal the effect of 17β-estradiol on an ER positive breast cancer cells. We observed that when treated with E2, cancer cells show an increase in the beta-oxidation rate as well as an increase in the cell proliferation rate. Moreover, the rate of *de novo* lipid synthesis in E2-treated cells was found to increase relative to untreated cancer cells. The elevated level of lipid consumption and lipid synthesis parameters in T47D cancer cells treated with E2 supports previous findings. The derived surge in cell proliferation rate matches previously published results, which relied on methylene blue staining [22, 23]. To support the fast cell proliferation, the cancer cells need to duplicate lipids to sustain the new cell membrane and organelles. Thus, the cells utilize the intermediate product of glycolysis (citrate) for *de novo* lipid synthesis rather than going through complete oxidative phosphorylation. Moreover, the binding of E2 to the estrogen receptor α (ERα) upregulates the expression of the glucose transporter and increases the influx of glucose molecules to support the biosynthesis. The adaption of cell glucose metabolism is confirmed by ORR measurements. Furthermore, the high metabolic activities inside cancer cells imply a huge demand for energy, which is met by combustion of the lipid storage.

These experiments proved the capability of combining NLOM and Michaelis-Menten modeling to quantify live cell metabolism with minimum interruption. Note that the data needed to be pre-processed to exclude lipid content in inactive cells to cover from complete β-oxidation inhibition to normal cell culture conditions.

## Author Contributions

<u>JH</u> Designed research; Performed research; Analyzed data; Wrote the manuscript

<u>NR</u> Designed research; Performed research; Developed kinetic models; Analyzed kinetic model with respect to data; Wrote the manuscript

<u>BTJ</u> Designed research; Wrote the manuscript

<u>EOP</u> Designed research; Wrote the manuscript

## Acknowledgement

This work was supported by National Institute of Biomedical Imaging and Bioengineering (P41EB015890), National Cancer Institute (R01CA142989 and 1R01CA195466-01), the NCI Chao Family Comprehensive Cancer Center (P30CA62203), UC HBCU program and the Arnold and Mabel Beckman Foundation.

